# Light might suppress both types of sound-evoked anti-predator flight in moths

**DOI:** 10.1101/727248

**Authors:** Theresa Hügel, Holger R. Goerlitz

**Author notes:** **corresponding authors:** Acoustic and Functional Ecology, Max Planck Institute for Ornithology, Eberhard-Gwinner-Str. 11, 82319 Seewiesen, Germany.

## Abstract

Urbanization exposes wild animals to increased levels of light, affecting particularly nocturnal animals. Artificial light at night might shift the balance of predator-prey interactions, for example of nocturnal echolocating bats and eared moths. Moths exposed to light show less last-ditch manoeuvres in response to attacking close-by bats. In contrast, the extent to which negative phonotaxis, moths’ first line of defence against distant bats, is affected by light is unclear. Here, we aimed to quantify the overall effect of light on both types of sound-evoked anti-predator flight, last-ditch manoeuvres and negative phonotaxis. We caught moths at two light traps, which were alternately equipped with loudspeakers that presented ultrasonic playbacks to simulate hunting bats. The light field was omnidirectional to attract moths equally from all directions. In contrast, the sound field was directional and thus, depending on the moth’s approach direction, elicited either only negative phonotaxis, or negative phonotaxis and last-ditch manoeuvres. We did not observe an effect of sound playback on the number of caught moths, suggesting that light might suppress both types of anti-predator flight, as either type would have caused a decline in the number of caught moths. As control, we confirmed that our playback was able to elicit evasive flight in moths in a dark flight room. Showing no effect of a treatment, however, is difficult. We discuss potential alternative explanations for our results, and call for further studies to investigate how light interferes with animal behaviour.

## INTRODUCTION

Light pollution by artificial light at night (ALAN) has increased substantially over the last decades (Fouquet 2006; Hölker *et al*. 2010; Falchi *et al*. 2016), adversely affecting plants and animals (Longcore and Rich 2004; Knop *et al*. 2017; Davies and Smyth 2018). The effects of light range from single individual’s orientation, reproduction and communication (Longcore and Rich 2004) to whole communities, for example, by shifting the balance of predator-prey interactions (Yurk and Trites 2000; Davies *et al*. 2013; Davies *et al*. 2014; Miller *et al*. 2017; Bailey *et al*. 2019; Russo *et al*. 2019). Echolocating bats and eared moths constitute a globally occurring predator-prey system of high ecological relevance (Boyles *et al*. 2011; Kunz *et al*. 2011; Kasso and Balakrishnan 2013; Van Toor *et al*. 2019). Their interactions take place in the darkness of the night and are exclusively mediated by sound. Echolocating bats hunt by emitting ultrasonic calls (Fenton 2003; Schnitzler *et al*. 2003), which eared moths can hear and react to with evasive flight (Roeder 1962; ter Hofstede and Ratcliffe 2016). Evasive flight likely consists of two stages: negative phonotaxis to fly away from distant bats (stage 1), and last-ditch evasive manoeuvres such as erratic flight or (power) dives to escape nearby attacking bats (stage 2). Corresponding to the differences in bat distance, negative phonotaxis is elicited at received sound pressure levels that are about 20 dB fainter than those that elicit last-ditch manoeuvres (Roeder 1962; Roeder 1964; Roeder 1967; Agee 1969).

Artificial light at night is of increasing concern for both bats and moths. While some bats may profit from exploiting prey accumulated at lights (Rydell 1992; Cravens *et al*. 2018), other species are negatively affected while foraging, commuting and roosting (Stone *et al*. 2009; Mathews *et al*. 2015; Stone *et al*. 2015; Straka *et al*. 2016; Straka *et al*. 2020). Moths are strongly attracted to light sources, leading to shortened foraging time (van Langevelde *et al*. 2017; Macgregor *et al*. 2019), disrupted navigation (Owens and Lewis 2018), reduced pollination (Macgregor *et al*. 2017), and population decline (van Langevelde *et al*. 2017; Wilson *et al*. 2018). Furthermore, light increases the predation risk of moths, for two reasons. The accumulations of moths around light sources attract bats, thereby increasing the predation pressure on moths (Rydell 1992; Cravens *et al*. 2018). In addition, light interferes with the moths’ sound-evoked anti-predator evasive flight response. In one set of studies, the sound-evoked evasive flight of moths was compared under lit and unlit conditions, showing that light reduces the evasive flight. Wakefield et al. (2015) showed that only 24% of moths performed last-ditch power-dives under LED illumination compared to 60% of moths in darkness; that is, the light inhibited last-ditch manoeuvres in 60% of the moths that would react in darkness. Similarly, Svensson & Rydell (1998) reported that moths within a radius of 1 m around a light source showed ~60% less last-ditch manoeuvres than moths in darkness (where 100% of moths reacted). Finally, Minnaar et al. (2015) reported the most extreme effect: in darkness, bat diet was best explained by a model that included evasive flight of moths. In contrast, with light, bat diet was best explained by a model that incorporated a 100% reduction in moth evasive flight, suggesting that the light inhibited both stages of evasive flight (negative phonotaxis and last-ditch manoeuvres).

In another set of studies, light exposure was kept constant while the sound received by the moth was manipulated. Those results showed that moths still exhibited some degree of evasive flight under illumination. Acharya & Fenton (1999) compared last-ditch manoeuvres in eared and deafened moths under illumination, showing that 48% of eared moths exhibited last-ditch manoeuvres when preyed on by bats, whereas deafened moths did not. Treat (1962) and Agee & Webb (1969) compared the number of caught moths at light traps with and without ultrasonic stimuli. Depending on sound stimulus and moth species, ultrasound playback reduced captures by 8-49 % (nine tympanate moth families with at least 39 caught individuals, Treat 1962) and by 51-86 % (in *Heliothis zea*, Noctuidae, Agee and Webb 1969) of the captures at the silent trap.

In summary, the first set of studies shows that light suppresses the sound-triggered evasive flight in 60-100% of the moths that would react in darkness. Contrasting this, the second set of studies shows that even in light, sound can still trigger evasive flight in 8-86% of the moths. Noteworthy, these studies either only reported effects of light on last-ditch manoeuvres (Svensson and Rydell 1998; Wakefield *et al*. 2015), or the results can be sufficiently explained by effects of light on last-ditch manoeuvres, as all moths had to fly through fields of high sound pressure level before entering the light trap (Treat 1962; Agee and Webb 1969). Only the modelling results of Minnaar et al. (2015) suggest that light completely suppresses both evasive flight responses under natural conditions. Therefore, while several lines of evidence suggest that last-ditch manoeuvres are suppressed by light pollution (with variable effect sizes), we lack a similar understanding of the effect of light on negative phonotaxis, and thus on the overall effect of light on evasive flight in moths. If the light-induced suppression of negative phonotaxis is as strong as for last-ditch manoeuvres, this will strongly affect the predator-prey interactions between bats and moths, because negative phonotaxis is elicited over larger distances and larger spatial volumes than last-ditch manoeuvres. Here, we advanced the light trap approach of Treat (1962) and Agee & Webb (1969) to investigate the effects of light on both stages of evasive flight, negative phonotaxis and last-ditch manoeuvres. We combined the omnidirectional light field of light traps (attracting moths equally from all directions) with a directional ultrasonic playback that should elicit different stages of evasive flight depending on each moth’s approach direction. As a moth approaches the light trap, its received light level will gradually increase independently from the approach direction, while its received sound pressure level will increase to different maximum values depending on the approach direction. Therefore, moths will be exposed to various combinations of light and sound pressure levels, covering a range of predator-prey scenarios (distant and close-by bats) and light levels (distant and close-by light sources) that a moth might encounter during the course of a night, allowing us to test the overall effect of light on both stages of evasive flight in moths. We compared moth captures at the light traps with and without acoustic playback, to measure the overall effect of light on the sound-evoked evasive flight. In line with Minaar et al. (2015) who suggest that light suppresses both stages of evasive flight, we predicted equal moth counts at both light traps, as either stage of evasive flight would cause a decline in the number of caught moths. In contrast, if negative phonotaxis (stage 1) was not suppressed or both stages were only partially suppressed, we predicted lower moth counts at the ultrasonic than the silent trap.

## METHODS

### Setup, study site, moth capture and measurement

We compared the number of moths caught at two light traps, one of which was equipped with a loudspeaker. We set up two equal light traps (Sylvania, blacklight, F15W/350BL-T8, USA, Fig. 1A); one next to a path in a forest (trap A) and the other one at 30 m distance in the forest (trap B), close to the Max Planck Institute for Ornithology, Seewiesen, Germany. Both traps hung freely at ~1.7 m above ground, radiating light at 360° in the horizontal plane (Fig. 1A). Between July 19 and August 16 2018, we collected data over 15 rainless nights, somewhat increasing the number of nights sampled in similar previous studies (2-12 nights (Treat 1962) and 6 nights (Agee and Webb 1969)). Each test night, we alternatingly equipped one of the two traps with two ultrasonic loudspeakers (ScanSpeak, Avisoft Bioacoustics, Glienicke, Germany); both broadcasting an ultrasonic stimulus (see below) to simulate echolocating bats. The two loudspeakers were fixed back to back facing in opposite directions and were mounted above the respective light trap at ~2 m above ground. Thus, each trap was associated for 7-8 nights with ultrasound simulating an echolocating bat. Each test night, lights and acoustic playback were turned on in the evening (between 20:30h and 23:40h) and turned off the next morning (between 7:00h and 9:40h). We emptied the traps each morning and counted all moths of the three ear-possessing families Noctuidae, Geometridae and Erebidae. We measured each individuals’ body length along the middorsal line (from the head to the end of the abdomen), to correct for the fact that larger moths have more sensitive hearing than smaller moths (Surlykke *et al*. 1999; ter Hofstede *et al*. 2013). For those individuals whose body length could not be measured (e.g., due to missing abdomen, N = 137, 15.5%), we either used the mean value of the species or, if this was not possible (N = 1), the mean value of the family. For statistical analysis, we binned individuals into six categories of body length (1.0 to 3.0 cm, bin width 0.5 cm).

**Figure 1:**
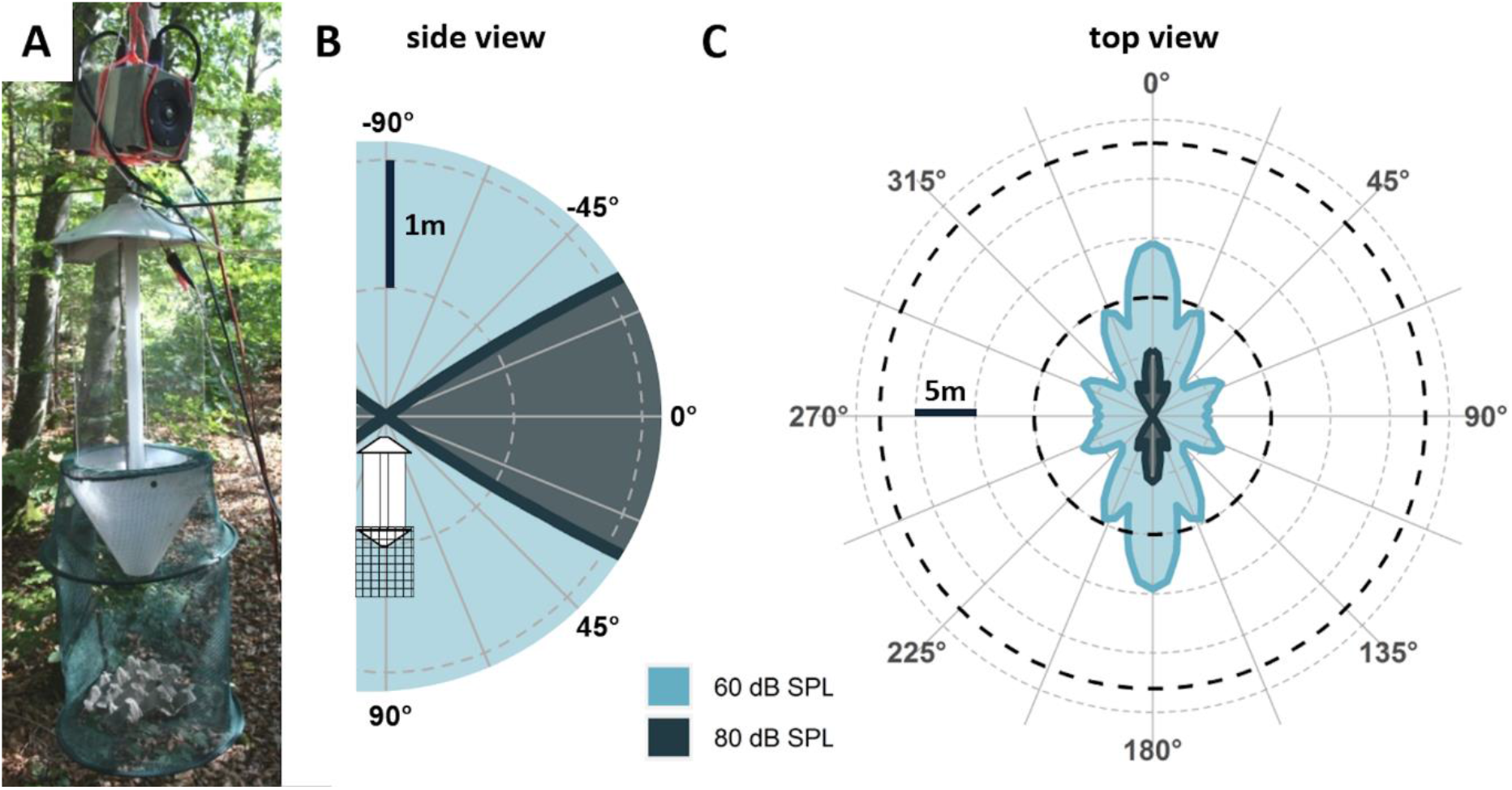
**(A) Experimental setup**. The photo shows one of the light traps with two loudspeakers attached above the trap, both of which broadcast the acoustic stimulus in opposite directions. **(B) Side view of the biologically relevant sound field around the light trap with attached loudspeakers**. Coloured areas indicate areas with minimum sound pressure levels of 60 and 80 dB SPL re. 20 μPa, which are biologically relevant acoustic thresholds for eliciting negative phonotaxis and last-ditch manoeuvres, respectively, in eared moths. **(C) Top view of the biologically relevant sound field (coloured areas) and light field (dashed lines)**. Dashed lines indicate the distance over which 5% of released noctuid (10 m) and geometrid (23 m) moths are recaptured at the light trap, respectively (Merckx and Slade 2014), which we used as first approximation of the range where light might interact with the moths’ sound-evoked anti-predator flight.

### Ultrasonic playback stimulus design and evaluation

We simulated predation pressure by echolocating bats by repeatedly playing a short bat-like ultrasonic pure tone pulse. Pulse frequency was 35 kHz, matching most moths’ best hearing threshold around 20-50 kHz (Noctuidae: ter Hofstede *et al*. 2013; Nakano *et al*. 2015; Erebidae: ter Hofstede *et al*. 2008; Pyralidae: Skals and Surlykke 2000; Geometridae: Rydell *et al*. 1997; Surlykke *et al*. 1997; Sphingidae/Drepanidae: Nakano *et al*. 2015). Pulse duration was 10 ms including 2 ms linear rise and fall times, corresponding to the calls of European open space bats (Obrist *et al*. 2004; Skiba 2014) and optimizing information transfer to the moths (Gordon and ter Hofstede 2018). Pulse interval was 100 ms, matching the call interval of searching bats (60 - 200 ms, Holderied and von Helversen 2003; Skiba 2014). On-axis sound pressure level (SPL) was 98 dB SPL re. 20 μPa RMS at 1 m distance (see below for a detailed description of the sound field). This stimulus was presented continuously in a loop throughout the night via the loudspeakers using Avisoft-RECORDER software (Avisoft Bioacoustics), a sound card with amplifier (Avisoft UltraSoundGate 116, Avisoft Bioacoustics) and a laptop computer.

To test the effect of our acoustic stimulus in darkness, without the potentially suppressing influence of light, we exposed free-flying moths in a dark flight room (5.3 m × 3.5 × 3 m^3^) to the same acoustic stimulus. We caught moths at trap A over the course of four nights and tested them within 30 hours after capture. We placed moths on the ground of the flight room and recorded the flight paths of upward-flying moths with an IR-sensitive camera (Sony HDR-CX560, Sony, Tokio, Japan) under IR illumination (850 nm, Mini IR Illuminator TV6700, ABUS Security-Center, Wetter, Germany). Using the same audio equipment as described above, we manually started the sound presentation when a moth flew in front of the speaker. We subsequently categorized the video-recorded flight paths as “reaction” when the flight direction, level of erraticness or both changed with stimulus onset (for examples, see supplementary video); as “no reaction” when we did not observe those changes; or as “ambiguous” when we could not clearly assign the flight path to one of the two previous categories.

### Overlap of sound and light field

The range and geometry of the presented light and sound fields differed. While the light was emitted omnidirectionally in the horizontal plane, the sound field was directional (Fig. 1). We estimated the biologically relevant range for attracting moths by light and the biologically relevant sound fields for triggering moths’ evasive flight based on literature values and our own measurements.

Light traps can attract released moths over up to 80 m distance, yet recaptures dramatically decrease beyond 15 m and depend on family (Truxa and Fiedler 2012; Merckx and Slade 2014). Family-specific models estimated the 5%-recapture rate at a distance of 10±6 m (mean ± SEM) for Noctuidae and 23±12 m for Geometridae (Merckx and Slade 2014). Note, however, that these distances were obtained with a different light source than ours (6W actinic vs. 15W blacklight in our case), and that the distance over which light attracts moths must not be equivalent to the distance over which light interferes with evasive flight. We still used these family-specific distances as first approximation for the biologically relevant light fields where light might interact with the moths’ sound-evoked anti-predator flight (Fig. 1B, dashed lines).

To estimate the effect of the playback, we measured the sound pressure level (SPL) of the pulse playback in front of the loudspeaker (on-axis) in steps of 5° up to 90° off-axis (for details, see SI). The on-axis source level at 1 ms distance was 97 dB SPL re. 20 μPa RMS, i.e., within the lower range of call levels emitted by free-flying bats (100-120 dB peSPL @ 1 m (Holderied and von Helversen 2003; Goerlitz *et al*. 2020), with the corresponding RMS-SPL-levels being ~3-7 dB lower than the peSPL-levels (Seibert *et al*. 2015; Lewanzik and Goerlitz 2017). With increasing off-axis angle, the source level became fainter by up to ~30 dB at 45°, resulting in a minimum playback level of 70 dB SPL RMS @ 1m. We then calculated the angle-dependent distances around the loudspeaker where the playback would reach biologically relevant levels of 60 and 80 dB SPL RMS. We chose 60 and 80 dB SPL as approximate thresholds for eliciting negative phonotaxis and last-ditch manoeuvres, respectively, based on several lines of evidence. Negative phonotaxis and last-ditch manoeuvres are likely triggered at levels just above the thresholds of the moths’ auditory receptor neurons A1 and A2, respectively (Roeder 1974; Madsen and Miller 1987; Gordon and ter Hofstede 2018). The best thresholds of the A1 neuron are at about 35-55 dB SPL, and of the A2 neuron at about ~52-72 dB SPL (Madsen and Miller 1987; Waters and Jones 1996; Surlykke 2003; ter Hofstede *et al*. 2013; ter Hofstede and Ratcliffe 2016; Gordon and ter Hofstede 2018). Behavioural thresholds in moths are largely unknown, but those that are known tend to be about 10 dB higher than neuronal thresholds (reviewed in Lewanzik and Goerlitz 2017). We thus defined 60 and 80 dB SPL RMS as thresholds that will likely elicit negative phonotaxis and last-ditch manoeuvres in most moth species, respectively, and calculated their isolines of constant sound pressure levels. SPL-Isolines varied with the angle around the loudspeaker, ranging from 3.6 to 14.6 m for 60 dB SPL RMS, and from 0 to 5.6 m for 80 dB SPL RMS (Fig. 1B). In summary, we thus presented a highly directional sound field in an omnidirectional light field. Thus, moths that approached the light trap in the on-axis direction of the loudspeaker experienced gradually increasing SPLs sufficiently high to first elicit negative phonotaxis and later last-ditch manoeuvres. In contrast, moths that approached from the side (off-axis to the loudspeaker) experienced gradually increasing SPLs that were only sufficiently high to elicit negative phonotaxis, but not last-ditch manoeuvres.

### Statistical analysis

To test for an effect of light on the moths’ evasive flight, we fitted linear models to the logarithmized moth count data as a function of the fixed effects *playback*, *trap*, *moth family*, and *moth body length*, and the interactions of *playback* and *moth family*, and of *playback* and *moth body length*.

Even though both model types could be fitted similarly well, we preferred a linear model with logarithmized count data over a negative bionomial model because the linear model enabled us to perform detailed power analysis. Although we have repeated measures over 15 nights, we did not include date as a random factor as it only explained a minor proportion of the variance in the logarithmized data.

We defined the moth family Noctuidae as intercept, as it had the largest sample size and thus was the most reliable reference. To test which factors significantly contributed to the model fit, we conducted backwards model reduction (Lewis *et al*. 2011). Hence, the full model was successively reduced, by stepwise removing factors, starting with the factor having the highest p-value of the t-statistics provided by the model summary. We compared models with likelihood ratio tests using a F-statistic and AICs. None of the interaction terms nor *trap* contributed significantly to model fit and were thus excluded. *Moth family* and *moth body length* contributed significantly to model fit (see results). Even though it did not contribute significantly to the model fit, we also kept *playback* as factor in the final model as this was the key parameter whose effect on the number of caught moths we aimed to analyse. Our final model thus included *playback*, *moth family* and *moth body length* as fixed factors, without any interactions.

We evaluated the power of our model for the effect sizes found in the field and in the flight room by a randomization approach of our real dataset. We first randomized the factor playback and then added an effect size as determined in the flight room or in the field to those logarithmized moth counts where the playback was “on”. We ran this simulation 10000 times for both effect sizes, and each time compared models (final model vs. final model without *playback*) with likelihood ratio tests using a F-statistic to test for a significant effect of *playback* on the model fit. The proportion of significant effects of *playback* per 10000 simulations equals our power to detect an effect of the tested effect size. All statistical analyses were conducted in R version 3.3.2 (R Foundation for Statistical Computing, Vienna, Austria) using the packages “lme4”(Bates *et al*. 2015), “MASS” (Venables and Ripley 2002), “blmeco” (Korner-Nievergelt *et al*. 2015), “DHARMa” (Hartig 2019), “car” (Fox 2019), “effectsize” (Kassambara 2019), and RNOmni” (McCaw 2018) for statistics and the packages “ggplot2” (Wickham 2020), “dplyr” (Wickham 2018), “cowplot” (Wilke 2019), and “ggthemer” (Arnold 2018) for data sorting and plotting. For further details, see R-script included in the SI.

## RESULTS

Of 33 moths tested in free-flight in the dark flight room, ten moths (30.3%) showed an evasive reaction in response to the acoustic stimulus. Twelve moths (36.4%) did not react, and eleven moths (33.3%) were categorized as “ambiguous”. The moths’ distance to the speaker at stimulus onset was about 1-2 m. Our acoustic stimulus thus was audible and elicited an evasive reaction in about one third of the tested moths. When considering the ambiguous reactions, the stimulus might even be audible to a larger proportion of up to two thirds of the moth population.

In the field experiment, we caught a total of 878 moths over 15 nights, with a median of ~23 moths caught per night and light trap (Fig. 2A), yet with large fluctuations between nights and smaller fluctuations between traps (7-116 moths per night and trap; Fig. 2B). We mostly caught moths of the family Noctuidae (80.9 %), followed by Geometridae (11.1%) and Erebidae (7.8%), and we caught more smaller than larger moths. Accordingly, both *moth family* and *moth body length* contributed significantly to our final model after stepwise model reduction (likelihood ratio tests: *moth family*, F(153,2) = 31.92, p < 0.001; *moth body length*, F_(153,3)_ = 34.43, p < 0.001).

**Figure 2:**
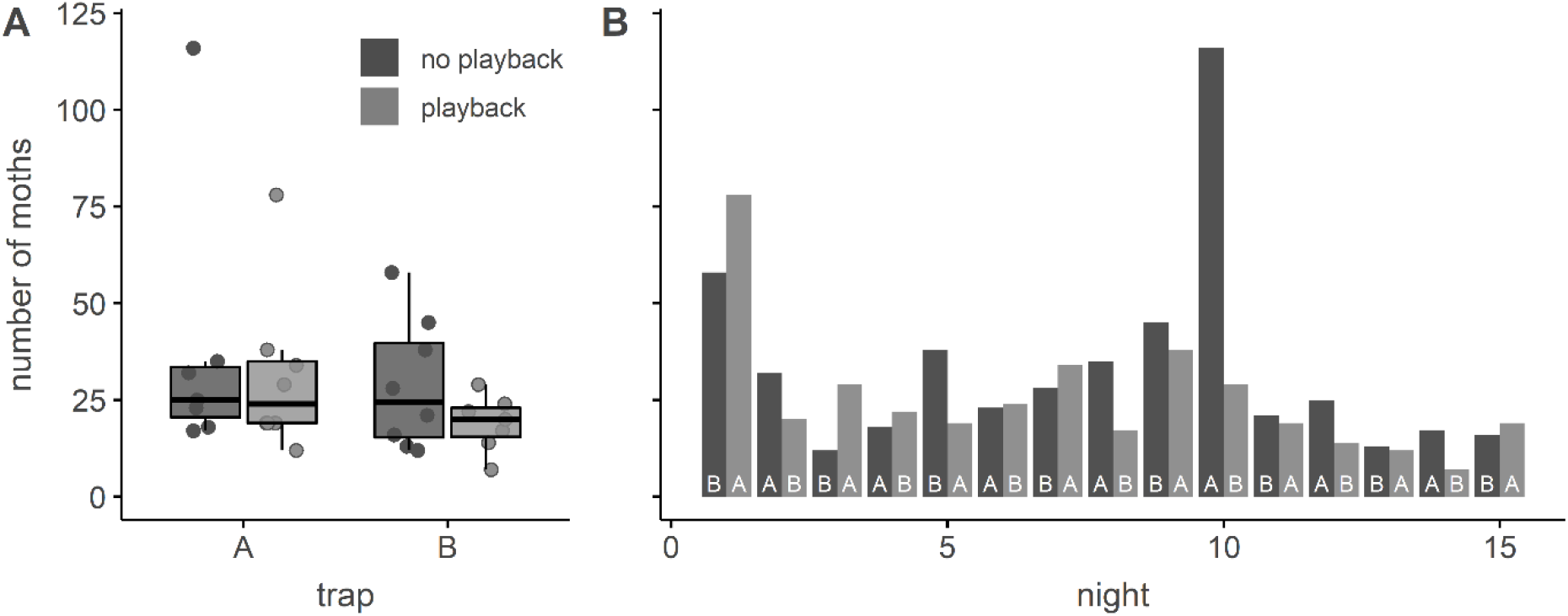
Number of caught moths per night and per playback treatment. **(A) Number of caught moths at both light traps, with and without playback**. Boxplots show median, quartiles and whiskers (up to 1.5 x inter-quartile range) of the daily counts (dots). **(B) Daily counts of caught moths per night and per trap for all 15 nights**. White letters at the base of the bars indicate the trap.

Although the total number of moths caught at trap A (506 moths) was 41% higher than the total number caught at trap B (365 moths), this was largely driven by one night (night 10: 115 vs 29 moths; Fig. 2B). Across all 15 nights, the nightly capture rate did not significantly differ between traps (likelihood ratio test for factor *trap*: F_(151,1)_ = 0.53, p = 0.468; Fig.2A).

We could not detect an effect of our acoustic playback on the nightly capture rates (Fig. 2A; modelled effect size of factor *playback* on logarithmized moth count at the playback trap relative to the silent trap: −0.06 (95% CI: −0.16 – + 0.04), corresponding to 86% (95% CI: 69 – 109%) capture rate at the playback trap relative to the silent trap; likelihood ratio test of factor *playback*, F_(152,1)_ = 1.53, p = 0.218).

We analysed the power of our experiment, both for the effect size observed in the field (−0.06) and the flight room. In the flight room, at least 30% of the moths reacted to our acoustic playback. Assuming that those reacting moths would not be caught in the light trap, the capture rate at the ultrasonic trap would be 70% relative to the silent trap, resulting in an expected effect size of the playback of −0.16 (in logarithmized moth counts). The power is the probability of rejecting the null hypothesis (i.e., obtaining a statistically significant result at a chosen significance level), given that the null hypothesis is false. Based on our dataset and sample size, our field experiment had a very low power of only 21% to detect an effect as small as the one observed in the field. In contrast, we had a sufficiently high power of 87% to detect an effect as large as the one observed in the flight room. This suggests that the lack of a significant effect of *playback* in the field might have been caused by a light-induced decrease of the effect size compared to the dark flight room – i.e., a light-induced suppression of sound-evoked anti-predator flight.

## DISCUSSION

Echolocating bats and eared insects are a textbook example of sound-mediated predator-prey interaction (ter Hofstede and Ratcliffe 2016). Increasing light pollution (Fouquet 2006; Hölker *et al*. 2010), however, severely impacts both bats and moths (e.g. Stone *et al*. 2009; Cravens *et al*. 2018; Macgregor *et al*. 2019), with potential cascading effects on their predator-prey-interactions, population dynamics and ecosystems (Minnaar *et al*. 2015; Russo *et al*. 2019). While good evidence exists that light reduces the sound-evoked last-ditch manoeuvres of eared moths, the effect of light on the moth’s first line of defence, negative phonotaxis, and thus the overall effect of light on moth anti-predator flight, is unclear. Here, we compared moth captures at two light traps. One trap was silent, while the other trap broadcast bat-like ultrasonic stimuli aimed to trigger last-ditch manoeuvres and negative phonotaxis. We did not find a significant reduction in the number of caught moths at the ultrasonic light trap. There is, however, a high bar for showing that a treatment (such as our ultrasonic playback) has no effect. The power of our experiment was too low to detect significant changes in moth count at an effect size as small as we observed in the field. In contrast, our field experiment had sufficient power to detect a potential effect of the playback with an effect size as large as observed in the dark flight room, if this effect had been present under lit field conditions. One conclusion thus is that the light suppressed both types of the moths’ sound-evoked anti-predator flight, negative phonotaxis and last-ditch manoeuvres.

There are, however, alternative explanations in addition to a light-induced suppression of anti-predator flight. The field and flight room experiments differed not only in the light level, but also in temporal (full night vs. short-term sound exposure) and spatial (variable distances between moth and loudspeaker vs close-range to the loudspeaker) parameters. In addition, physiological and behavioural states of the moths will likely differ between free-flying moths in the field and captured and released moths in the flight room.

The continuous ultrasonic playback over a full night might cause the moths to habituate to the playback. Habituation was previously suggested as an explanation for playback-independent capture rates of male *Helicoverpa zea* moths at pheromone traps (Gillam et al 2011), although it was not shown at the neuronal level in response to searching bats (Gordon and ter Hofstede 2018). However, although our playback was on throughout the night, the extent to which a given individual moth was exposed to the playback depends on the speed and shape of its own flight behaviour. Arguably, our experimental situation might be similar to the realistic case of moths around street lights that are attacked by circling close-by bats (Rydell 1992). Indeed, Treat (1962) and Agee & Webb (1969), whose studies our study was based on, also presented ultrasonic playbacks throughout the night and were able to detect differences between the silent and ultrasonic trap, arguing against habituation. We also believe that differences in stimulus design cannot explain the differences between experiments. Treat (1962) and Agee & Webb (1969) broadcast multiple stimuli varying in pulse rate, frequency, duration, and sound pressure level (ranges: 0.7-155 pulses/s; 12.5-200 kHz; 2-10 ms; SPL: ~60-100 dB SPL @ 1 m distance), all of which elicited varying degrees of evasive flight in eared moths. Our stimulus had acoustic properties within this range and did elicit evasive flight under dark control conditions, yet seems not to elicit sufficient evasive flight under lit conditions.

When assuming that light indeed suppressed the moths’ sound-evoked antipredator-flight in our experiment, the question arises why this was not the case in the similar light-trap experiments by Treat (1962) and Agee & Webb (1969). We propose that the differences in the geometry and overlap of light- and sound-fields might explain these contrasting results. In the previous setups (Treat 1962; Agee and Webb 1969), sound and light fields almost overlapped and were emitted within a relatively narrow angle. Before entering the light trap, approaching moths thus passed through high sound pressure levels that would likely elicit last-ditch manoeuvres. As both studies caught fewer moths in the ultrasonic trap than the silent traps, this suggests that the playback still elicited some anti-predator flight (likely last-ditch manoeuvres) despite the light. Specifically, for stimuli similar to ours, the relative capture rates between the ultrasonic and silent traps were 35% vs. 65% for noctuid moths and a 37.5 kHz tone pulses (Treat 1962), and 15% vs. 85% for two species of the families Noctuidae and Pyralidae and a 30 kHz tone pulses (Agee and Webb 1969). In contrast, our setup combined an omnidirectional light field with a directional sound field, thus exposing the moths to different sound pressure levels (SPL) depending on approach direction. When approaching the trap on-axis of the loudspeaker’s main axis, received SPLs increased from low to high, which should first elicit negative phonotaxis and later last-ditch manoeuvres. In contrast, when approaching the traps off-axis, SPLs remained so low to only elicit negative phonotaxis. As our capture rates did not differ among the light traps – indicating that the playback did not evoke anti-predator flight – we suggest that the light not only reduced last-ditch behaviour, but also the negative phonotaxis of eared moths. Even if the moths still exhibited some degree of last-ditch manoeuvre close to the trap (as shown by Treat 1962 and Agee and Webb 1969), these manoeuvres might have brought the moth into a position off-axis to the loudspeaker with low SPL (either to the side or below the loudspeaker’s main axis). From there, no further last-ditch manoeuvres would have been elicited due to the low SPL, while the light kept attracting the moth into the trap and suppressed negative phonotaxis. Our results are in line with those of Minnaar et al. (2015), who found that a model of escape behaviour in moths assuming 0% efficiency best explained bats’ diet under lit conditions. We therefore suggest that light suppresses not only last-ditch manoeuvres, as previously shown, but also negative phonotaxis. Further experiments need to validate this suggestion, by testing low and high source levels separately (either separated spatially or temporally), by increasing the number of nightly and spatial replicates, and by testing additional bat-like sounds for a variety of light and sound level combinations. In addition, tracking and quantification of the three-dimensional evasive flight of individual moths of different species will provide details insight into the variability of their evasive flight, and the effect of light on it.

In summary, our results underline the strong effect of light on eared moths, and suggest that both types of anti-predator flight are suppressed by light. It is important to note, though, that our study design tested the effect of light only indirectly, by testing the effect of sound on light-mediated moth captures, not the effect of light on sound-mediated evasive flight. In addition, showing no effect is difficult, and is complicated by the natural variability of moth behaviour (Hügel & Goerlitz 2019), potentially complicating simple answers. If increasing artificial light at night suppresses both negative phonotaxis and last-ditch manoeuvres, moths are not only unable to escape nearby predators, but also unable to avoid distant predators by flying away. Similarly, fish are attracted to lit areas, where they are “trapped” and preyed upon by seals (Yurk and Trites 2000) and other fish (Becker *et al*. 2013). The increasing levels of light pollution demand for further studies to understand the mechanism(s) of how light attracts animals and interferes with their behaviour.

## Supporting information

Supplementary Infromation

## Ethics

Permission to carry out fieldwork was provided by the relevant German authority (Landratsamt Starnberg, permit # 55.1-8646.NAT_02-8-41).

## Data accessibility

All data, R-scripts and a flight video are provided as electronic supplementary material.

## Author contributions

TH and HRG conceived of and designed the study. TH collected and analysed the data. TH and HRG wrote the manuscript. All authors gave final approval for publication and agree to be held accountable for the work performed therein.

## Acknowledgements

We thank the IMPRS for support, the participants of the Scientific Writing Course 2019 and two anonymous reviewers for comments on previous versions of the manuscript, and the Max Planck Institute for Ornithology Seewiesen for excellent infrastructure and support.

