## Supplementary Infromation for "Light might suppress both types of sound-evoked anti-predator flight in moths"

### Supplementary Information

#### Measurements and calculations for the sound fields of 60 and 80 dB SPL RMS

We measured the emitted sound pressure levels (SPL) at three different distances (1, 2, and 4 m; twice per distance) and at all angles from 0° (on-axis) to 90° off-axis in steps of 5° around the speaker (Fig. S2). Next, we corrected the measurements taken at 2 and 4 m distance for atmospheric attenuation (21.4°C, 42% rel. humidity) and geometric attenuation due to spherical spreading and back-calculated them to the SPLs at 1 m distance. Including the original measured values at 1 m distance, this resulted in 3-6 values for 1 m distance per angle (missing values due to clipping). We next calculated the mean of all values at 1 m distance per angle to obtain the angle-dependent sound pressure levels. These mean values were used as angle-depended source level of the loudspeakers to calculate the sound field around the loudspeaker (with atmospheric attenuation for 21.4°C and 42% rel. humidity and spherical spreading). Resulting sound fields can be seen in 1 B & C.

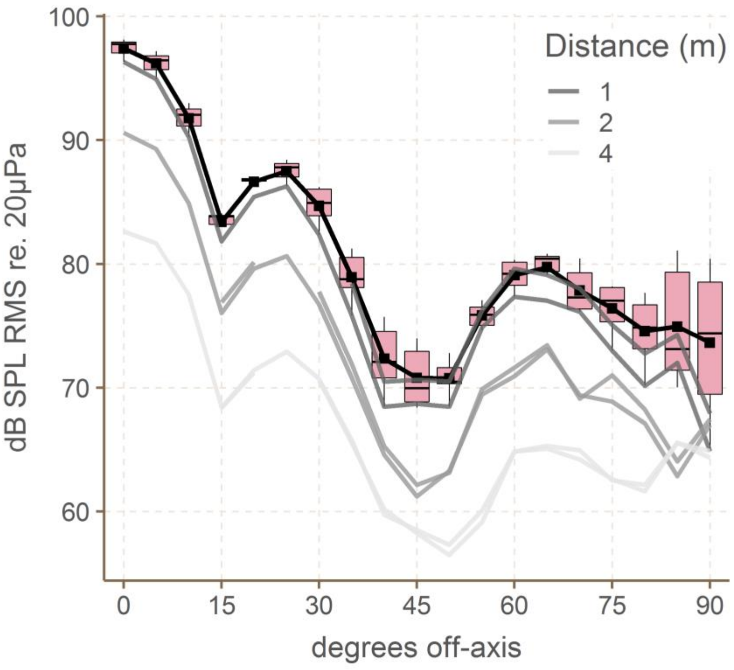


**Figure S2: Measurement of the sound pressure level (SPL) around the loudspeaker**. Measurements were taken at three distances (1,2 and 4 m) and angles from 0° to 90° in steps of 5°. Lines show the original measurements at all three distances, once measured from 0-90 degrees and once measured from 90-0 degrees. Missing values are due to clipping. Boxplots show the SPL at 1 m distance, combined from the original measurements at 1 m distance and back-calculated from the measurements taken at 2 and 4 m distance, indicating median, quartiles, whiskers (up to 1.5x the inter-quartile range beyond the quartiles). Black squares show means, which are connected by the black line.
